# DCP: a pipeline toolbox for diffusion connectome

**DOI:** 10.1101/2020.04.16.044453

**Authors:** Weijie Huang, Ni Shu

## Abstract

The brain structure network constructed by diffusion tensor imaging (DTI) reflects the anatomical connections between brain regions, so the brain structure network can quantitatively describes the anatomical connectivity pattern of the entire brain. The structure network based diffusion tensor imaging is widely used in scientific research. While a number of post processing packages have been developed, fully automated processing of DTI datasets on Windows Operating System remains challenging. Here, we developed a MATLAB toolbox named “Diffusion Connectome Pipeline” (DCP) for fully automated constructing brain structure network. The processing modules of a few developed packages, including Diffusion Toolkit, DiffusionKit, SPM and MRIcron, were employed in DCP. Using any number of raw DTI datasets from different subjects, in either DICOM or NIfTI format, DCP can automatically perform a series of steps to construct network. In addition, DCP has a friendly graphical user interface (GUI) running on the Windows Operating System, allowing the user to be interactive and to adjust the input/output settings, as well as the processing parameters. As an open-source package, DCP is freely available at https://www.nitrc.org/projects/dcp. This novel toolbox is expected to substantially simplify the image processing of DTI datasets and facilitate human brain structural connectome studies.

## Introduction

The human brain is one of the most complex system in nature. Different functional areas interact and coordinate with each other so that they can complete a variety of basic physiological functions and advanced cognitive functions. However the brain neurobiological mechanisms remain so far unclear. In recent years, researchers have proposed the using of network models and topological measures based on graph theory to study the structure and function of the brain (Bassett and Bullmore, 2006; Bullmore and Sporns, 2009). By building a network model of the brain, a large number of studies have consistently found that the brain network has small world attributes (Bassett and Bullmore, 2006) and modular structure (Chen et al., 2008; Meunier et al., 2009). This implies that the human brain has evolved into a complex but efficient nervous system in order to achieve efficient and synchronous interaction between brain regions and to differentiate and integrate functions (Sporns and Zwi, 2004; Achard and Bullmore, 2007).

The human brain contains 10^11^ neurons and 10^15^ synapses. Building brain networks from the micro level of the neurons is not only technically impossible, but contains too much redundant information from a global point of view, so the study of brain network at present is dividing the brain into different areas and construct a network basing the connections between the brain areas. According to different technical means, there are two main categories of network which are function networks and structure networks.

The connections of structure networks reflect the anatomical connections between the brain regions, so the research on structure network can quantitate the anatomical connectivity pattern of the entire brain. It is very important for understanding the structure underlying the brain function. From the physiological meaning, the topological distance between nodes is closely related to physiological distance between brain areas in structure network. Considering the long distance connections will consume more energy and material, the adjacent brain regions are more likely connected together in the structure networks to meet minimize energy consumption and wiring volume.

The advent of brain imaging techniques has made possible the study of living human brain structure networks. The vivo brain structure network can be constructed based on structure magnetic resonance imaging (MRI) or diffusion tensor imaging (DTI) technology. Because different modality images reflect different brain information, the construction of the structure network is different. Study on the structure MRI based on the cortical thickness correlation in different brain regions thought there are connections between brain regions with high correlation. The brain network is constructed indirectly. The structure network constructed by He and his colleague based cortical thickness correlation of 124 normal controls was most representative (He et al., 2007). The study of diffusion MRI, through the fiber tracking algorithm, directly reconstructed the anatomical connections between different brain regions (Hagmann et al., 2008; Iturria-Medina et al., 2008; Gong et al., 2009), and thus obtained the anatomical connection structure network.

However, the complex preprocessing steps of DTI greatly limit its application. Nowadays, although there are many software to process DTI data, they all have some limitations. Such as FMRIB Software Library (FSL) (Smith et al., 2004), it can only be run on Linux Operating System. For users unfamiliar with Linux, it is not convenient to use; DTI-studio (Jiang et al., 2006) and DiffusionKit (Xie et al., 2016) are run on the Windows Operating System, but they are operated step by step which is same as FSL. When there are a large number of data, not only the operation is complicated but also is easy to make a mistake. And those software can’t construct the structure network based on the whole brain fiber imaging. Although PANDA (Cui et al., 2013) has batch processing function, but it can only run on Linux Operating System which most of the researchers aren’t accustomed to using.

In order to simplify the automatic processing of neuroimaging data, some scholars have now developed a cross platform library. The library provides a framework for automatic processing of neuroimaging data, but it requires the users to customize their own module and the parameters used by the program. This is not friendly to no encoding experience scientific researchers.

Therefore, we have developed a MATLAB toolkit based on Windows Operating System for building DTI networks named Diffusion Connectome Pipeline (DCP). Firstly, it provides a friendly graphical user interface, and the users only need to use the mouse to set the parameters which are used in the whole process and click the button with label ‘RUN’. The DCP will process every participant’s data automatically, and generate DTI network of every participant. And the default parameters we provide are currently recognized as classic processing parameters. When the program is running, the user can accurately get which step the program is running through the monitor window. After the program is finished, a folder of quality control is generated, and the images of registration results of each subject are saved for quality checking.

## Materials and methods

DCP was developed based on MATLAB in Windows Operating System. Lots of toolkits from SPM, MRIcron, Diffusion Toolkit and DiffusionKit were called by DCP. The pipeline processing in DCP will be described here followed by an introduction to the realization of pipelines.

### Overview of functionality of DCP

We show all procedures of DCP in **Figure 1** and it includes four steps: (1) preprocessing; (2) tractography; (3) generating parcellation; and (4) constructing matrix.

**Figure 1.**
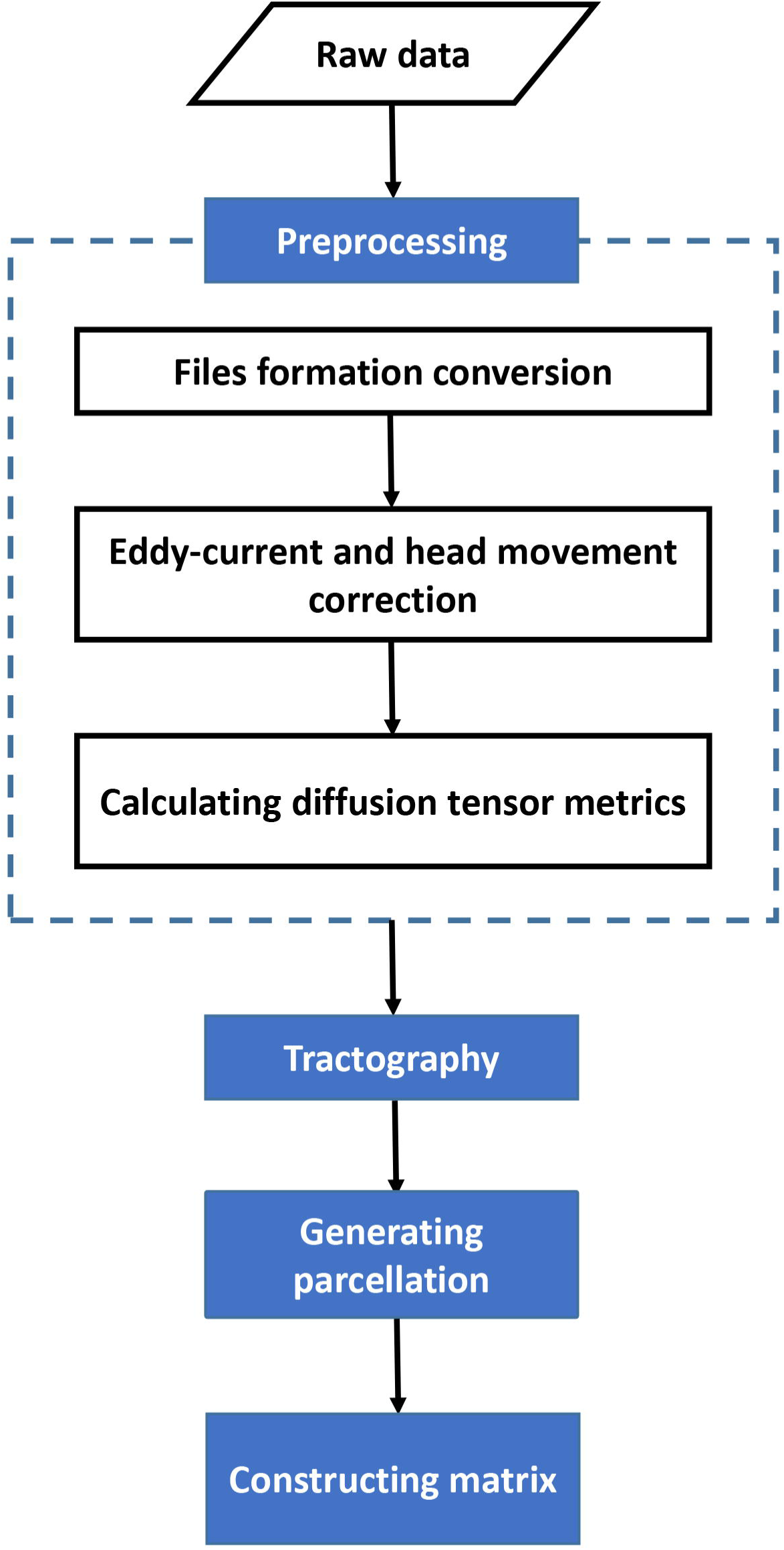
Main procedure for pipeline processing of DTI datasets in DCP. The procedure includes four parts: (1) preprocessing; (2) tractography; (3) generating parcellation; and (4) constructing networks.

#### Preprocessing

In this section, to construct DTI networks, DCP allows researchers to perform several preprocessing steps of DTI data that are commonly used in the community and it includes three steps: (1) files formation conversion; (2) Eddy-current and head movement correction; (3) Calculating diffusion tensor metrics.

#### Files formation conversion

The DICOM data is a format output from most MRI scanners. DCP can only process NIFTI data. So the first step of processing data is converting DICOM files into NIFTI files. This conversion is achieved in DCP by calling dcm2nii tool in the MRIcroN software (http://people.cas.sc.edu/rorden/mricron/index.html) or SPM (http://www.fil.ion.ucl.ac.uk/spm/).

#### Eddy-current and head movement correction

The distortion of DTI would be induced by eddy-current and simple head-motion during scanning. This distortion can be corrected by registering the diffusion weighted images to the b0 image with an affine transformation. DCP achieve this transformation by calling bneddy command in DiffusionKit (Xie et al., 2016).

#### Calculating diffusion tensor metrics

This step calculates tensor matrix as well as diffusion tensor metrics including fractional anisotropy (FA), apparent diffusion coefficient (ADC), etc voxel wisely. The dti_recon command in diffusion toolkit was applied.

#### Tractography

In this step, DCP uses deterministic fiber tracking. Every stage work of fiber tracking algorithm can be described as follows: (1) select an interested seed zone; (2) from the seed point, find the next voxel along with the fiber direction ; (3) repeat (2) until you reach the end of your head, or meet a certain stop conditions (FA below a threshold value, or the fiber direction mutation); (4) from the seed point, repeat (3) along with the opposite direction of the previous track to construct the other half fiber; (5) for each seed point in the seed zone, repeat (2)-(4) step. DCP achieve this by applying dti_tracker command in diffusion toolkit.

#### Generating parcellation

The DCP divides the entire brain into multiple regions using a prior gray matter (GM) atlas, where each region represents a network node (Bullmore and Sporns, 2009). Nevertheless, the prior atlases need to be transformed to the native DTI space of each individual from the standard space. In order to solve this problem, DCP uses the co-register, normalize and deformation toolbox in SPM. Specially, the individual structural image (i.e, T1-weighted) is co-registered to its corresponding individual b0 image using the co-register toolbox in SPM and uses the b0 image to estimate brain mask by the bet command of MRIcroN. Then use the mask to remove the skull in individual structure image which is co-registered b0 image space. The individual structure image which is co-registered b0 image space is then non-linearly registered to the ICBM152 template with the normalize toolbox in SPM. An inverse warping transformation from the standard space to the native DTI space can be obtained by the resultant transformations in these two steps. Applying this inverse transformation to warp inversely prior atlases in the standard space back to individual native space. Currently There are two well-defined atlases: the Automated Anatomical Labeling(AAL) (Tzourio-Mazoyer et al., 2002) atlas and a atlas which is randomly subdivided into 1024 regions with equal size (Zalesky et al., 2010) were provided by DCP. And of course, other customized atlases can be imported into DCP to define the network nodes.

#### Constructing matrix

The schematic flowchart of network construction is demonstrated in **Figure 2**. For every pair of brain nodes/regions defined above, they are considered structurally connected if there are at least one fiber streamline with two end-points that were located in these two regions (Shu et al., 2011; Zalesky et al., 2011; Bai et al., 2012). Based on the linking fibers, DCP can calculate three basic weighted matrices: fiber number-weighted matrix (FN), FA-weighted matrix (FA), and length-weighted matrix (Len). Each row or column represents a brain region/node in the matrices. The values of the elements FN(i, j), FA(i, j), and Len (i, j) represent the fiber number, averaged FA and averaged length of linking fibers between node i and node j, respectively. The resultant matrices were saved as MATLAB data files which can be directly used for topological analysis with graph theoretic approaches and text files for checking.

**Figure 2.**
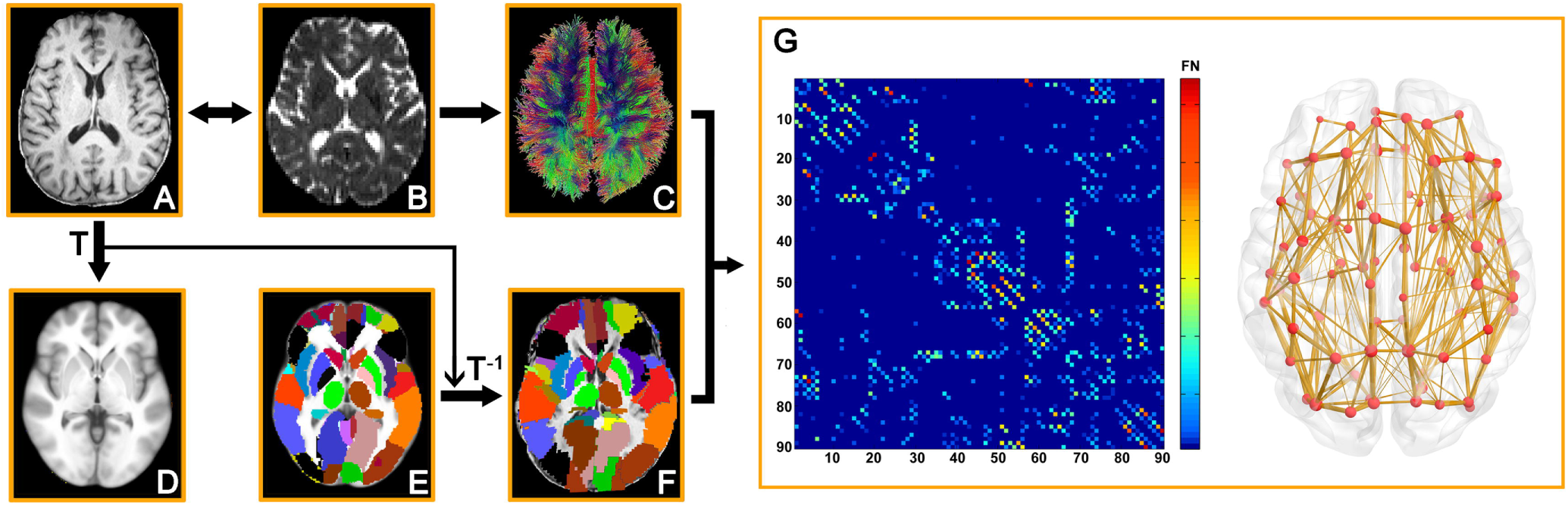
Flowchart for constructing anatomical brain networks using diffusion tractography in DCP. (A) Structure image(T1-weight) in diffusion space. (B)b0 image. (C) White matter tracks constructed using deterministic tractography. (D) Structure image in standard space. (E) Parcellation of grey matter in standard space. Each color represents a node in network. (F) Parcellation of grey matter in diffusion space. (G) The network matrix weighted by fiber number.

### Testing the group differences on structure network metrics by using DCP

#### Subjects

Fifty-one aMCI patients were recruited from the Memory Clinic of the Neurology Department, XuanWu Hospital, Capital Medical University, Beijing, China and Fifty-one demographically matched healthy controls (HCs) were recruited from communities using advertisements. All of the participants were Han Chinese and right-handed. The following inclusion criteria of HCs were used: (1) no complaints of memory loss or related disorders causing cognitive impairment; (2) a CDR score of 0; (3) no severe visual or auditory impairment. Participants were excluded if they met any of the following conditions: (1) a history of stroke, addictions, neurological/psychiatric diseases, or treatments that would affect cognitive function; (2) major depression (Hamilton Depression Rating Scale score > 24 points); (3) other central nervous system diseases that may cause cognitive impairment, such as brain tumors, Parkinson’s disease, encephalitis, or epilepsy; (4) cognitive impairment caused by traumatic brain injury; (5) other systemic diseases, such as thyroid dysfunction, severe anemia, syphilis, or HIV; (6) large vessel disease (such as cortical and/or subcortical infarcts and watershed infarcts); or (7) the diseases with WM hyperintensity or WM lesions (such as normal pressure hydrocephalus and multiple sclerosis).

This study was approved by the Institutional Review Board of XuanWu Hospital, Capital Medical University. Written informed consent was obtained from each participant. The main demographic and clinical information of all participants is presented in **Table 1**.

**Table 1.** Demographic of all participants

| | aMCI (n = 51) | NC (n = 51) | T/$\chi^2$ value | P value |
| --- | --- | --- | --- | --- |
| Age (years) | 64.1±9.5 (42-82) | 62.2±9.1 (43-83) | 0.99 | 0.32<sup>a</sup> |
| Gender (M/F) | 22/29 | 18/33 | 0.66 | 0.42<sup>b</sup> |
| Education (years) | 9.6±4.1 (0-18) | 11.2±4.6 (0-19) | -1.93 | 0.06<sup>a</sup> |

Data are presented as the Mean ± SD (range). Abbreviations: aMCI, amnesic mild cognitive impairment; NC, normal control; <sup>a</sup>The P value and T value were obtained by two-sample t-test. <sup>b</sup>The P value and $\chi^2$ value were obtained by the $\chi^2$ test.

#### MRI acquisition

All participants were scanned using a Siemens Trio 3.0 T MRI scanner at XuanWu Hospital, Capital Medical University, China. Participants lay still with their heads fixed by straps and foam to minimize movement. The magnetization prepared rapid gradient echo (MPRAGE) sequence was used to obtain the T1-weighted images with the following parameters: repetition time (TR), 1,900 ms; echo time (TE), 2.2 ms; flip angle, 9°; acquisition matrix, 256×224; field of view (FOV), 256×224 mm2; slice thickness, 1 mm; no gap; 176 sagittal slices; resolution, 1 mm3 isotropic; and average, 1. The diffusion tensor imaging (DTI) data were acquired using a single-shot echo-planar imaging-based sequence with the following parameters: TR, 11,000 ms; TE, 98 ms; flip angle, 90°; acquisition matrix, 128×116; FOV, 256×232 mm2; slice thickness, 2 mm; no gap; 60 axial slices; resolution, 2 mm3 isotropic; and average, 3. Thirty non-linear diffusion weighting directions with b = 1,000 s/mm2 and one b0 image without diffusion weighting (b = 0 s/mm2) were obtained. Additionally, an axial FLAIR/T2-weighted and resting-state functional MRI images were also collected for each participant. All images were reviewed and the leukoencephalopathy and vascular comorbidity was evaluated by an experienced neuroradiologist.

#### Image processing

The whole pipeline procedure of DCP was run on all DTI data to construct anatomy network. We set seed number to 1, turning angle threshold to 45, lower FA threshold to 0.2, higher FA threshold to 1 in tractography and used AAL atlas and H1024 atlas to define nodes. And constructed FN-weighted network, FA-weighted network and Len-weighted network. Finally we used GRETNA (Wang et al., 2015) to calculate graph metrics.

#### Network topology

We applied Graph theoretical approaches to characterize the topology of brain networks which are derived from neuroimaging data (Bullmore and Sporns, 2009). Here we pay attention to two graph metrics: global efficiency and local efficiency. Global efficiency was defined as the average of the inverse of the “harmonic mean” of the characteristic path length, which represented the ability of network to process information parallel (Latora and Marchiori, 2001). The local efficiency of network which quantify the fault-tolerant ability of network is the average of nodal local efficiency that is defined as the global efficiency of the subgraph composed by its nearest neighbors (Latora and Marchiori, 2001).

#### Statistical analysis

For all network attributes, we used two sample t-test to calculate the between-group differences in their mean values. The subjects’ age, sex and education were regressed as covariates with general linear model (GLM) and p < 0.01 was considered as significant.

## Results

### An integrated MATLAB toolbox: DCP

In this study, We developed a MATLAV toolbox called DCP which is an open-source package and is freely available at https://www.nitrc.org/projects/dcp. The online discussion forum (https://www.nitrc.org/forum/?group_id=1180) and a mailing list (https://www.nitrc.org/mail/?group_id=1180) have been registered for DCP, and we will provide technical supports and updates constantly.

The DCP can construct structure networks based on DTI. Not only can it be batch-processed, it can also be executed separately by a single step in the program (e.g., DICOM conversion, tractography, generating parcellation and constructing matrix). Particularly, DCP has a very friendly GUI, with which users can perform various interactions with the embedded functions, e.g., setting inputs or outputs and configuring the processing parameters. In addition, the user can get the status of the program running in real time from the DCP’s GUI. After the program finished, the native space parcellation will be saved as a PNG picutre, for the user to check whether there is any error in the process of generating the parcellation.

### Resultant files of DCP

Firstly, DCP will recognize the modality of MRI and rename the files containing DTI image as DTI_* (* represented 1, 2, 3 and so on), and rename the files containing T1 images as T1_* (* represented 1, 2, 3 and so on). Then for each subject, DCP will generate three folders as listed in **Table 3**. Specifically, the DTI_DATA folder consists of result files of preprocess and tracktography. The files in the PARCELLATION includes the result files of generating parcellation. The folder named MATRIX containes the all matrix files. At last, users can set a output file which will be generated in the parent folder of input folder if users didn’t set. It containes a folder named QC which consisted of quality control files of all subjects and a MAT format file which contains all the subjects’ networks.

**Table 2.** Group differences on WM connectivity metrics between aMCI patients and NC.

| Network | Metric | MCI (n = 51) | NC (n = 51) | T value | P value |
| --- | --- | --- | --- | --- | --- |
| FA, AAL | Global efficiency | 0.22±0.02<br>(0.17-0.25) | 0.22±0.01<br>(0.20-0.26) | 2.58 | 0.011 |
|  | Local efficiency | 0.30±0.02<br>(0.24-0.35) | 0.31±0.01<br>(0.27-0.34) | 2.76 | 0.007 |
| FA, H1024 | Global efficiency | 0.09±0.01<br>(0.04-0.12) | 0.10±0.01<br>(0.07-0.12) | 3.50 | <0.001 |
|  | Local efficiency | 0.16±0.02<br>(0.11-0.21) | 0.18±0.02<br>(0.13-0.21) | 3.37 | 0.011 |
| FN, AAL | Global efficiency | 13.28±2.61<br>(8.11-17.76) | 15.21±2.27<br>(9.60-19.48) | 3.82 | <0.001 |
|  | Local efficiency | 20.38±3.70<br>(13.23-28.40) | 22.82±3.08<br>(15.37-29.84) | 3.45 | <0.001 |
| FN, H1024 | Global efficiency | 0.93±0.19<br>(0.49-1.30) | 1.05±0.17<br>(0.67-1.46) | 3.32 | 0.001 |
|  | Local efficiency | 2.10±0.32<br>(1.47-2.86) | 2.27±0.32<br>(1.52-2.96) | 2.69 | 0.009 |

Data are presented as the Mean ± SD (range). FA means the fractional anisotropy-weighted network; FN means the fiber number-weighted network. AAL means using the Automated Anatomical Labeling atlas to define the nodes. H1024 means using the atlas which is randomly subdivided into 1024 regions with equal size to define the nodes.

**Table 3.** Folders produced by DCP

| Folder name | Files |
| --- | --- |
| DTI_* | Raw DTI files |
| DTI_DATA | Resultant files of preprocess and tracktography |
| PARCELLATION | Resultant files of generating parcellation |
| MATRIX | Resultant files of constructing network |
| TI_* | Raw T1 files |

### The group differences on structure metrics by using DCP

As shown in **Table 2**, in FA weighted network constructed with AAL template, the local efficiency and global efficiency decreased significantly in aMCI patients compare with NCs. The global efficiency and local efficiency of FA weighted network constructed by H1024 template in aMCI patients were smaller than NCs. And in FN weighted network constructed by AAL template or H1024 template, significant decreased global efficiency and local efficiency were observed in the aMCI patients compared with HCs. These results are coincident with previous studies (Shu et al., 2012).

## Discussion

In this study, we developed a MATLAB toolbox named DCP to process DTI data automatically. The toolbox has two key advantages. The first advantage is it is run in the Windows Operating System not the Linux Operating System which the PANDA and FSL was run in. The second advantage is that it fully automates all the processing steps of DTI datasets for any number of subjects.

A fully automatic pipeline naturally makes the data processing more efficient, at the same time reducing potential mistakes due to human intervention. Although users can use MIPAV (McAuliffe et al., 2001), JIST (Lucas et al., 2010), Nipype (Gorgolewski et al., 2011), or LONI (Dinov et al., 2009) for DTI data processing, but these require prior knowledge on pipeline design and programming skills related to these package. In addition, knowledge of the details of DTI data processing is also required, which will be another challenge for end users. In order to provider end users with an easy to use tools, The DCP was developed which make it possible to construct anatomic networks automatically and immediately for any number of subjects.

Notably, processing procedures across existing DTI packages was slightly different, and might overlook some important processing steps (Jones et al., 2013). These issues have been well discussed by a few recent articles (Jones and Cercignani, 2010; Jones et al., 2013). The processing pipelines of DCP have tried to follow the best practice as much as possible. For example, the adjustment of diffusion gradient directions after eddy-current correction, which has been frequently missed (Leemans and Jones, 2009; Jones et al., 2013), has been included in the DCP pipeline. And when co-register the T1 image to DTI space we used the b0 image which contained less noise instead of FA image. Then used the b0 image without skull as mask to remove the non-brain area in T1 image which was beneficial to the next normalize. In future versions, DCP will keep being updated to include processing steps of the best practice at the moment.

Finally, the advanced users can select the desired options for each processing step with friendly GUI (**Figure 3**) of DCP. Users need to change some settings that fit their data for the best quality of processing. Users can also replace the reference data, e.g., image templates for normalization or prior atlases for node definition to their customized data to make it possible for processing DTI data of non-human (e.g., primate) brains.

**Figure 3.**
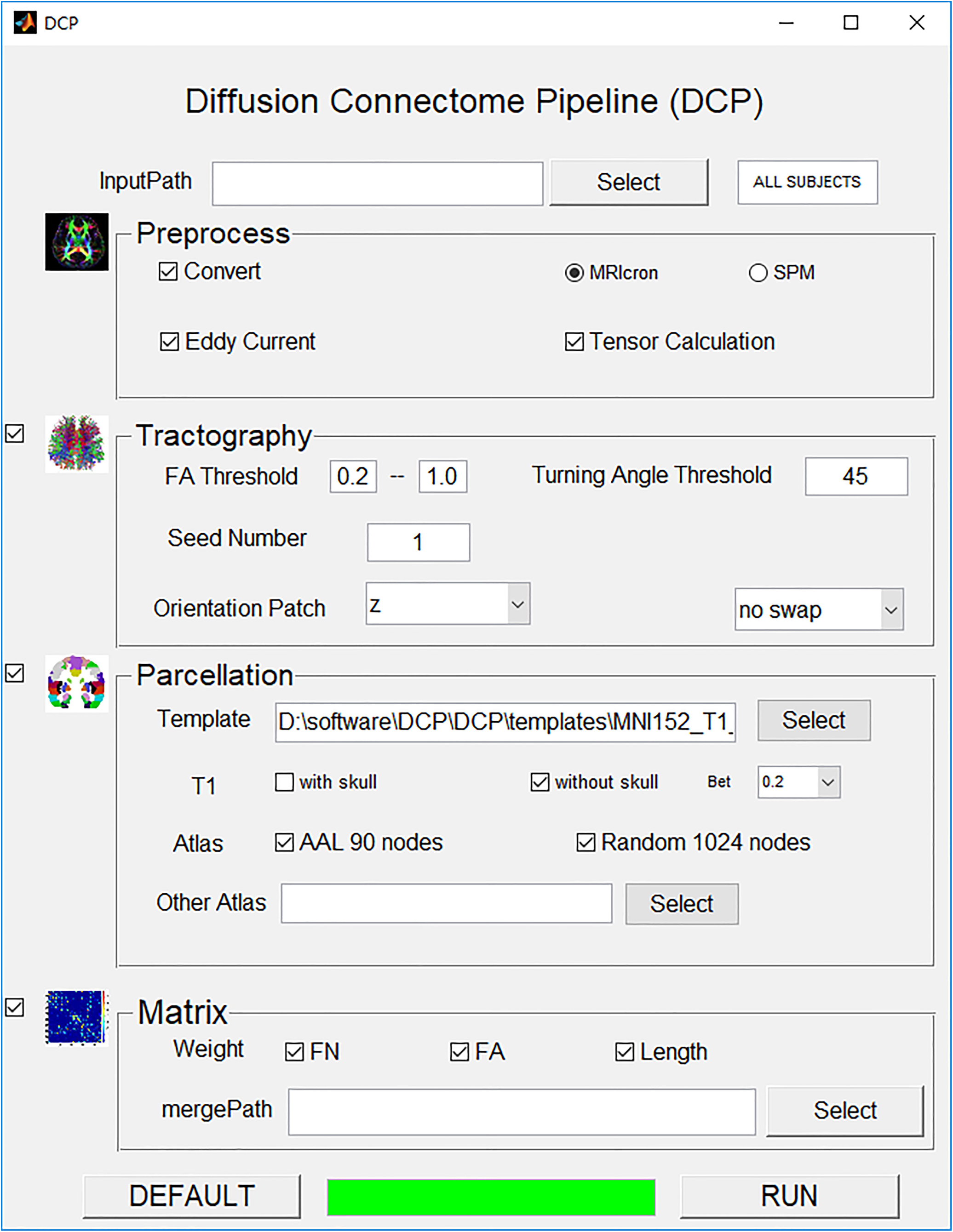
A snapshot of the GUI of DCP. The GUI allows for inputting raw DTI datasets and configuring processing parameters and monitoring the progress of data processing in real-time.

In the present study, we applied DCP to construct WM network based aMCI and normal control’s DTI images. And test the network topology metrics betweenaMCI and NC. We found that the NC had higher global efficiency in FA weighted network, FN weighted network and length weighted network. These results are consistent with previous findings (Zhao et al., 2017).

Since 2005, Olaf Sporns put forward the concept of connectome (Sporns et al., 2005), more and more attention has been paid to the research of brain network., and DTI has been taken as the main technical means tobuild structural brain network (Behrens and Sporns, 2012). This will lead to a large number of DTI datasets in the foreseeable future (http://humanconnectome.org/). To process these connectome dataset, DCP has unique advantages, as it can handle the large number of datasets automatically. Therefore, DCP will contribute to the study of the human connectome in the near future.

In summary, DCP can construct three types weighted WM network (fiber number weighted network, FA weighted network and length weighted network) from raw DICOM files conveniently with friendly GUI. And it is update easily because of its extendable design framework.

